# An alphavirus replicon-based vaccine expressing a stabilized Spike antigen induces sterile immunity and prevents transmission of SARS-CoV-2 between cats

**DOI:** 10.1101/2021.04.01.436305

**Authors:** Martijn A. Langereis, Ken Stachura, Suzan Miller, Angela M. Bosco-Lauth, Irina C. Albulescu, Airn E. Hartwig, Stephanie M. Porter, Judith Stammen-Vogelzangs, Mark Mogler, Frank J.M. van Kuppeveld, Berend-Jan Bosch, Paul Vermeij, Ad de Groof, Richard A. Bowen, Randy Davis, Zach Xu, Ian Tarpey

## Abstract

Early in the global SARS-CoV-2 pandemic concerns were raised regarding infection of other animal hosts and whether these could play a significant role in the viral epidemiology. Infection of animals could be detrimental by causing clinical disease but also of concern if they become a viral reservoir allowing further mutations, plus having the potential to infect other animals or humans. The first reported animals to be infected both under experimental conditions and from anecdotal field evidence were cats described in China early in 2020. Given the concerns this finding raised and the close contacts between humans and cats, we aimed to determine whether a vaccine candidate could be developed that was suitable for use in multiple susceptible animal species and whether this vaccine could reduce infection of cats in addition to preventing spread to other cats.

Here we report that a Replicon Particle (RP) vaccine based on Venezuelan equine encephalitis virus (VEEV), known to be safe and efficacious for use in a variety of animals, expressing a stabilised Spike antigen, could induce neutralising antibody titers in guinea pigs and cats. After two intramuscular vaccinations, virus neutralising antibodies were detected in the respiratory tract of the guinea pigs and a cell mediated immune response was induced. The design of the SARS-CoV-2 antigen was shown to be critical in developing a strong neutralising antibody response. Vaccination of cats was able to induce a serum neutralising antibody response which lasted for the course of the experiment. Interestingly, in contrast to control animals, infectious virus could not be detected in oropharyngeal or nasal swabs of vaccinated cats after challenge. Moreover, the challenged control cats spread the virus to in-contact cats whereas the vaccinated cats did not transmit virus. The results show that the RP vaccine induces sterile immunity preventing SARS-CoV-2 infection and transmission. This data suggests that this RP vaccine could be a multi-species vaccine useful for preventing spread to and between other animals should that approach be required.

## Introduction

SARS-CoV-2 is an extremely contagious respiratory coronavirus that emerged in China in late 2019 and has since spread globally causing the on-going coronavirus disease 2019 (COVID-19) pandemic. Coronaviruses are enveloped, single stranded, non-segmented, positive sense RNA viruses that encode sixteen non-structural proteins and four structural proteins. The structural Spike protein is the major determinant of host cell tropism by binding to the angiotensin-converting enzyme 2 (ACE2) on cells, a type I integral membrane protein that plays an important role in human vascular health. Using the ACE2 receptor to gain entry to cells in the upper respiratory tract (URT) SARS-CoV-2 infection of humans has manifested itself in a wide range of clinical outcomes from asymptomatic to very severe respiratory infections which in some situations are complicated by immunological dysfunction causing COVID-19 with over 2.7 million fatalities to date. As ACE2 receptors that are highly similar to the human receptor are also present on the cells of a number of other animals it is important to understand whether those potential hosts can play any role in disease spread. Since the first human infections it has been shown that cats, dogs, ferrets, hamsters and mink can be readily infected either in laboratory studies or via natural transmission [1–5]. The role these susceptible animals play in the human epidemiology is unclear though two-way transmission between mink and humans has been demonstrated leading to the culling of all animals in mink farms [3]. It is therefore of upmost importance to understand the role of animals in the spread of this virus, especially with regard to their potential to act as a viral reservoir, and the possibility to develop important viral variants which thereby influence the overall epidemiology.

The importance of cats in the epidemiology of COVID-19 has yet to be fully established, though there are a significant number of reports of cats testing positive for SARS-CoV-2, mostly in association with human infections in the same household. The first published report demonstrating that cats could be experimentally infected also showed virus transmission to in-contact cats [1]. Whilst the infected cats did not demonstrate overt clinical disease, significant respiratory lesions were detected post-mortem especially in younger cats. In subsequent experimental trials no clinical disease was observed in challenged cats but prolonged shed of virus and spread to in contact cats was again detected [4][5]. In addition to these experimental infection studies, there have been numerous reports of domestic cats testing positive for SARS-CoV-2 with less than a quarter showing signs of disease and no severe presentations as reported in humans [6]. Although a number of these cases were associated with the presence of a confirmed SARS-CoV-2 infected owner, this was not always the case [7] and natural infection between domestic cats has not been ruled out. Serological surveys of cats in China [7], USA [8], France [9] and Italy [10] have demonstrated that a high proportion of cats tested positive for SARS-CoV-2, including feral animals with no known history of ownership.

Although concerns regarding feline infections have significantly reduced, the initial reports that a large number of cats were being abandoned by owners [11] led to key opinion leaders releasing statements regarding the low risk of human infection from cats [6]. Furthermore, owners testing positive for SARS-CoV-2 have been advised to distance themselves from their cats in an attempt to prevent transmission, and SARS-CoV-2 infection of cats is now reportable to the OIE [12]. Recently, with the rise of new variants, there are also reports that these variants may have altered host tropism [13] and possibly also different pathogenesis [14]. For these reasons it is important to further study the epidemiology of SARS-CoV-2 in cats and whether the possibility exists of the feline population becoming a natural reservoir for the virus.

A large number of human vaccines are now in development against SARS-CoV-2, with more than five approved for use in various regions globally. The types of vaccine include adjuvanted expressed SARS-CoV-2 Spike protein, adjuvanted whole SARS-CoV-2 virus vaccines, mRNA encoding SARS-CoV-2 Spike, and recombinant viral vector vaccines expressing the SARS-CoV-2 Spike protein [15]. Early reports indicate that these vaccines have good safety profiles and have greatly reduced both the number and severity of infections. Other important considerations for the long term success of these vaccines include, the immunological correlates of the protection induced, the vaccination scheme required to induce an appropriate duration of immunity, the cost and production scale of these vaccines required for the global population and whether any antibody dependent enhancement is detected as has been seen on rare occasions with other coronavirus vaccine candidates [16]. The use of coronavirus vaccines in the veterinary industry is well established with a variety of vaccines against infectious bronchitis virus (IBV), bovine coronavirus (BCV), porcine epidemic diarrhoea virus (PEDV), feline infectious peritonitis (FIP) and canine coronavirus (CCV) being used broadly for many decades. As an interesting parallel to SARS-CoV-2 infection, IBV is transmitted via the respiratory route initially causing an upper respiratory infection in chickens followed by systemic disease which, depending on the strain, can involve the kidney, reproductive organs, or intestinal tract. IBV has evolved into an enormous number of variant strains globally and it is important to note that many different serotypes of IBV are present [17]. These serotypes are sufficiently antigenically distinct that most require unique serotype specific vaccines to control disease.

The licensure of IBV vaccines requires not only the demonstration of protection from clinical disease but also a highly significant reduction of virus replication in the trachea. Given its respiratory route of transmission and the requirements to significantly reduce virus present in the respiratory tract, the most effective vaccines are live attenuated viruses which are delivered by mucosal application, by spray or in drinking water. Local delivery of live attenuated IBV vaccines induces relatively short-lived local mucosal IgA neutralising antibody responses in addition to systemic IgA and IgG antibody responses [18–20]. However, the complete mechanism of protection is unclear and is likely to involve cell mediated immunity specific for other proteins besides Spike. In longer lived birds, responses are boosted by the parenteral delivery of adjuvanted inactivated whole virus vaccines which extends the duration of immunity significantly and is likely to boost the immune response that has been primed by the mucosally delivered vaccine. Interestingly the induction of virus neutralising serological antibodies against the Spike protein by parenteral vaccination routes has previously been shown to provide a low level of protection against respiratory IBV challenge [21–23]. It is therefore of particular interest to determine how well the human SARS-CoV-2 vaccine candidates, most of which are designed to be administered parenterally, are able to control upper respiratory tract infection and virus spread, in addition to preventing clinical disease. Initial indications are that these vaccines are successfully reducing human to human spread though the mechanisms involved require further investigation.

In the controlled experiments in felines it has been shown that the SARS-CoV-2 virus can readily infect cats and although in most cases no or only a mild disease was detected, the cats can shed the virus for prolonged periods infecting other cats [4]. Prevention or limitation of virus replication in infected cats, and thereby reducing direct contact transmission in cats, would be useful in limiting the establishment of a reservoir in the feline population and also potentially limit the development or selection of viral mutants in these species. In order to investigate prevention of transmission between cats we tested whether a vaccine could protect cats from SARS-CoV-2 challenge and also prevent virus spread between cats under controlled conditions.

For this purpose, we utilised an Alphavirus based Replicon technology derived from the attenuated TC-83 strain of Venezuelan equine encephalitis virus (VEEV) to express the SARS-CoV-2 Spike protein. Replicon technology has been tested in numerous species (including humans) [24,25] and this VEEV based replicon has been shown to be safe in cats reducing the clinical effects and shed of Feline Calicivirus which causes an acute respiratory tract infection (Authors unpublished observations). Furthermore, good responses have been detected in chickens, dogs, horses, pigs and cattle with a variety of antigen targets [24]. The replicon forms the basis of the Sequivity^®^ RNA Particle vaccine platform which is currently licensed in the US for multiple swine applications. In this system the foreign gene of interest, in this case SARS-CoV-2 Spike, is inserted in place of VEEV structural genes generating a self-amplifying RNA capable of expressing the gene of interest upon introduction into cells. The self-amplifying replicon RNA directs the translation of large amounts of protein in transfected cells, reaching levels as high as 15–20% of total cell protein [26]. As the replicon RNA does not contain any of the VEEV structural genes, the RNA is propagation-defective. The replicon RNA can be packaged into replicon particles (RP) by supplying the VEEV structural genes *in trans* in the form of promoterless capsid and glycoprotein helper RNAs and, when the helper and replicon RNAs are combined and co-transfected into cells, the replicon RNA is efficiently packaged into single-cycle, propagation-defective RP which are used in the vaccine formulation [27]. RP vaccines have been shown to induce both innate and adaptive immune responses including virus neutralising antibodies and T cell responses [24]. Of significant importance is that this system can be employed very rapidly with materials sufficient availability for deployment of vaccine within weeks.

The structural conformation and localisation of the SARS-CoV-2 Spike protein has been found to be important for induction of a protective immune response [28–30]. We therefore generated two RP vaccine candidates producing either the wildtype Spike antigen (Spike^WT^) or an optimised Spike protein antigen (Spike^Opt^). These candidate vaccines were assessed for Spike protein expression and stability *in vitro*, and then for the induction of a mucosal and serological antibody response and T cell stimulation following subcutaneous vaccination of guinea pigs. Following these experiments, the vaccine demonstrating the most optimal characteristics was tested by subcutaneous vaccination for the induction of a serological response in cats followed by a mucosal SARS-CoV-2 challenge and monitoring for clinical signs, shed of virus orally or nasally and transmission to in-contact non-vaccinated cats.

## Materials and Methods

### Animals and husbandry

Female SPF guinea pigs (Dunkin Hartley) were obtained from Envigo at a minimum weight of 350 grams, randomly allocated to experimental groups and individually marked using color coded tags. Baseline clinical observations were documented throughout the study period. Domestic short hair male and female SPF cats were obtained from Marshall BioResources (Waverly, NY), identified by microchip and randomly allocated to experimental groups. Baseline clinical observations including body temperatures were documented throughout the study period.

### Generation of SARS-CoV-2 Spike gene replicon particle (RP) vaccines

The VEEV replicon vectors used to produce either the SARS-CoV-2 Spike^WT^ or Spike^Opt^ gene were constructed as previously described [31] with the following modifications. The TC-83-derived replicon vector “pVEK” was digested with restriction enzymes *AscI* and *PacI* to create the vector “pVHV”.

The Spike^WT^ gene sequence from SARS-CoV-2, strain 2019-nCoV/USA-WI1/2020 (GenBank accession MT039887), and the Spike^Opt^ derivative possessing the R^68^2A/R^683^A (ΔFCS) K^986^P/V^987^P (2P) substitutions and replacement of SARS-CoV-2 Spike residues 1212-1273 for residues 463-511 of VSV glycoprotein (GeneBank accession YP_009505325, were codon-optimized for expression in cat and synthesized with flanking *AscI* and *PacI* sites (ATUM, Newark, CA). The synthetic genes and pVHV vector were each digested with *AscI* and *PacI* enzymes and ligated to create vectors “pVHV-SARS-CoV-2-Spike^WT^” and “pVHV-SARS-CoV-2-Spike^Opt^”. Plasmid batches were sequenced to confirm the correct vector and insert identities.

Production of TC-83 RNA replicon particles (RP) was conducted similarly to methods previously described [32]. Briefly, pVHV-SARS-CoV-2-Spike^WT^ and pVHV-SARS-CoV-2-Spike^Opt^ replicon vector DNA and helper DNA plasmids were linearized with *NotI* restriction enzyme prior to *in vitro* transcription using RiboMAX™ Express T7 RNA polymerase and cap analog (Promega, Madison, WI). Importantly, the helper RNAs used in the production lack the VEE subgenomic promoter sequence, as previously described [27]. Purified RNA for the replicon and helper components were combined and mixed with a suspension of Vero cells, electroporated in 4 mm cuvettes, and returned to serum-free culture media. Following overnight incubation, alphavirus RNA replicon particles were purified from the cells and media by passing the suspension through a depth filter, washing with phosphate buffered saline containing 5% sucrose (w/v), and finally eluting the retained RP with 400 mM NaCl + 5% sucrose (w/v) buffer or 200 mM Na_2_SO_4_ + 5% sucrose (w/v) buffer. Eluted RP were passed through a 0.22 micron membrane filter and dispensed into aliquots for storage prior to assay and lyophilisation. A control vaccine was also prepared expressing green fluorescent protein. The titers of functional RP-Spike vaccines were determined by immunofluorescence assay on infected Vero cell monolayers following lyophilisation in a stabiliser containing sucrose, NZ Amine and DMEM and storage at 2-8°C. Briefly, the vaccine was serially diluted and added to a Vero cell monolayer culture in 96-well plates and incubated at 37°C for 18-24 hours. After incubation, the cells were fixed and stained with the primary antibody (anti-VEEV nsp2 monoclonal antibody) followed by a FITC conjugated anti-murine IgG secondary antibody. RNA particles were quantified by counting all positive, fluorescent stained cells in 2 wells per dilution using the Biotek^®^ Cytation™ 5 Imaging Reader.

### Placebo control vaccine

The placebo vaccine consisted of RNA Particles expressing the green fluorescent protein (GFP) assayed, lyophilized and stored at 2-8°C as described above. Following use, each of the test vaccines were titrated to confirm the vaccination dose.

### Guinea pig study

SPF guinea pigs with a minimum weight of 350 grams were randomly divided over the non-vaccinated control group, RP-Spike^WT^ vaccine group, and RP-Spike^Opt^ vaccine group (n=6 per group). One week after placement, animals remained either non-vaccinated or received a prime vaccination of 1 × 10^7^ RP dose intramuscularly (0.1 ml in each leg muscle). Three weeks after prime vaccination animals received a booster vaccination of 1 × 10^7^ RP dose intramuscular (0.1 ml in each leg muscle). Six weeks after the booster vaccination animals received a second booster vaccination and 7 days later animals were sacrificed. Terminal blood was taken for LST and trachea were carefully dissected without causing bleedings. Mucus was taken from the inside of the trachea using a swab, taken up in 1 ml of phosphate buffered saline and used to determine mucosal antibody titers. At the day of booster vaccination, and with 2-week interval until 6 weeks after boost vaccination, clotted blood was taken using cardiac-puncture and serum was used to determine systemic antibody titers.

### Surrogate VN assay Guinea pig sera

The SARS-CoV-2 Surrogate Virus Neutralization Test Kit from GenScript (REF: L00847) was used according to manufacturer’s instructions. Briefly, sera were diluted in sample dilution buffer, mixed 1:1 with HRP-RBD, and incubated 30 minutes at 37°C. Next, samples were put in a 96-well plate containing ACE2 receptor coated on the surface and incubated 15 minutes at 37°C. Unbound HRP-RBD was washed away and remaining HRP was visualized using TMB substrate and measured at OD450.

### ELISA for estimating anti RBD and SED antibody titers in sera

Purified SARS-CoV-2 RBD and SED (Spike ectodomain) were diluted in DPBS (without Ca and Mg, Lonza, 17-512F) and coated onto 96-well plates (MaxiSorp - ThermoFisher or High binding - Greiner Bio-one) using 10nM (10 pmols/mL), and incubated overnight at 4°C. Next morning plates were washed with an ELISA plate washer (ImmunoWash 1575, BioRad) using 0.25 mL wash solution/well (DPBS, 0.05% Tween 20) three times, then blocked with 250 μL blocking solution (5% milk - Protifar, Nutricia, 0.1% Tween 20 in DPBS) for 2 hours at RT (room temperature). Afterwards the blocking solution was discarded, 4-fold serial dilutions of the sera (prepared in blocking solution, in duplicates or triplicates) were added to the corresponding wells and incubated for 1h at RT. Each plate contained positive control (guinea pig sera diluted to obtain an OD450 of ~2) and negative control wells. Plates were washed again 3 times before being incubated with the HRP-containing antibody – Goat anti-Guinea pig (IgG-HRPO, Jackson Lab 106-035-003, 1:8000) for 1 hour at RT. The last wash steps were performed, followed by incubation for 10 minutes at RT with 100 μL/well Super Sensitive TMB (Surmodics, TMBS-1000-01). Reactions were stopped by adding 100 μL/well of 12.5% H2SO4 (Millipore, 1.00716.1000). Absorbance at 450 nm was measured within 30 minutes with an ELx808 Biotek plate reader.

### T-cell stimulation test (LST)

Blood was collected and lymphocytes were isolated using Sepmate tube (Stemcell) containing Histopaque 1083 according to manufacturer’s instructions. Briefly, K3-EDTA blood was diluted 1:2 in RPMI-1640 medium and pelleted for 10 minutes at 1.200 × *g*. Cells in the top layer of the tubes were collected, put in a clean tube containing RPMI-1640 and pellet for 7 minutes at 400 x g. Cells were washed once with RPMI-1640 medium and pelleted for 7 minutes at 400 × *g*. Cell concentrations were counted and 1 × 10^7^ cells were stained with CFSE for 20 minutes at 37°C. Cells were washed once with RPMI-1640 and from each animal 5 × 10^5^ cells were stimulated with either medium, ConA (10 μg/ml), or purified SARS-CoV-2 S1 antigen (5, 2.5, 1.25, 0.62, 0.31, or 0.15 μg/ml) in duplicate. Three days after stimulation, cell proliferation was measured using the FACS-Verse.

### SARS-CoV-2 challenge virus and cell culture

SARS-CoV-2 strain USA-WA1/2020 (GenBank: QHO60594.1) was isolated from an oropharyngeal swab from a patient with a respiratory illness who had returned from travel to an affected region of China and developed clinical disease (COVID-19) in January 2020 in Washington, USA. The virus was propagated for one passage on Vero cells. To determine the virus titer, serial dilutions of virus were made on Vero cells and plaque forming units quantified by counterstaining with a secondary overlay containing Neutral Red at 24 hours and visualization after 48 hours of incubation.

### Feline Serology

Serological responses to SARS-CoV-2 were studied using an *in-vitro* plaque reduction neutralisation test (PRNT). Briefly serum was inactivated at 56°C for 30 minutes, serial dilutions of cat serum were prepared and incubated with 100 pfu of SARS-CoV-2 for one hour at 37°C. The virus serum mixtures were then plated onto Vero cells and the number of plaques read by counterstaining with a secondary overlay containing Neutral Red at 24 hours and visualization after 48 hours. Antibody titers were determined as the reciprocal of the highest dilution in which ≥90% of virus was neutralised.

### Efficacy test

Two groups of ten 11-week-old SPF cats were formed and housed separately; one group was vaccinated with 5 × 10^7^ RP-Spike^Opt^ by the subcutaneous route (0.5ml per dose) with the other group receiving the same dose of RP-gfp. After three weeks each group received the same treatment. Twenty-five days following the second vaccination the cats were challenged as previously described [4] though using both the intranasal and oral routes with 3.1 × 10^5^ pfu of SARS-CoV-2 under light sedation. A further two groups of five SPF cats that were neither vaccinated nor challenged were used as sentinels by co-housing with each group 1 day post-challenge. All animals were observed daily for 10 days following challenge for clinical signs indicative of SARS-CoV-2 infection. Clinical signs checked included depression, dyspnea, nasal discharge, ocular discharge, cough, conjunctivitis, and/or sneezing. Body temperatures were recorded on study days 1-11 post-challenge/post-mingling.

### Oropharyngeal swabs

Oropharyngeal swabs for virus isolation were collected from the challenged cats on study days 1 to 7 post-challenge, the swabs were placed in Tris-buffered MEM containing 1% bovine serum albumin containing gentamycin, penicillin, streptomycin and amphotericin B (BA-1 media). To assess contact spread swabs were also collected from the contact sentinels into transport media on study days 2-8 post-challenge. The samples were frozen at −50°C until testing.

### Nasal washes

Nasal wash samples for virus isolation were collected days 1, 2, 3, 5 and 7 post-challenge as previously described [4] by instilling 1 ml of BA-1 media into the nares of cats and collecting nasal discharge in a petri dish. To assess contact nasal washes were also collected from the contact sentinels on days 2, 3, 4, 6, and 8 post-challenge. The samples were frozen at −50°C until testing.

### Blood samples

Blood samples were taken for sera prior to and 3 weeks post primary vaccination. In addition, blood samples were taken prior to and 14 days post challenge.

### Virus re-isolation

All oropharyngeal swabs and nasal washes were tested for virus re-isolation as previously described [4]. Confluent monolayers of Vero E6 cells in 6 well plates were washed once with PBS and seeded with 100ul of serial ten-fold dilutions of swab/wash samples, incubated at 37°C for one hour then overlaid with 0.5% agarose in MEM containing 2% FBS. A second overlay containing neutral red dye was added 24 hours later and plaques counted at 48 hours. Viral titers were recorded as log_10_ pfu/ml.

### Statistical analyses

A two-tailed T test was used to compare serological responses. P-values of less than 0.05 were considered to be significant.

## Results

### SARS-CoV-2 Spike antigen design

The recent SARS-CoV-2 vaccine efforts have unambiguously shown that stabilizing the pre-fusion form of the Spike protein enhances immunogenicity of the antigen in the mRNA and vector-based vaccines [33]. Also, we have shown for the IBV Spike protein that replacing the transmembrane (TM) and C-terminal domain (CTD) for its counterparts of vesicular stomatitis virus (VSV) glycoprotein enhanced cell-surface localization in vitro and immunogenicity *in vivo* (Authors unpublished observations). Therefore, we designed an optimized Spike antigen (Spike^Opt^) comprising an inactivated furin cleavage site (FCS) as well as the introduction of the double-proline (2P) substitution to stabilize the pre-fusion conformation of the antigen. Additionally, the TM and CTD of SARS-CoV-2 Spike has been replaced by the similar domains of the VSV glycoprotein (Figure 1).

**Figure 1:**
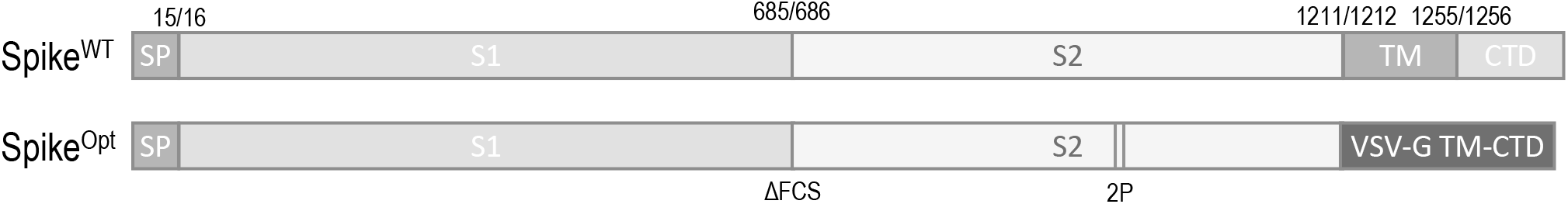
Schematic representation of the wildtype SARS-CoV-2 Spike antigen (Spike^WT^) and the stabilized SARS-CoV-2 Spike antigen (Spike^Opt^). Different Spike protein domains are indicated by different grey shadings. The furin cleavage site mutation (ΔFCS, R^682^A/R^683^A), 2P substitutions (K^986^P/V^987^P) and TM-CTD replacements are shown.

### Immunogenicity study of RP-Spike vaccine candidates in guinea pigs

Immunogenicity of the Spike^WT^ and Spike^Opt^ antigens was assessed in a guinea pig model in which the VEEV RP vector vaccines were given intramuscularly (Figure 2A). After prime vaccination all animals showed seroconversion as assessed by a commercially available surrogate VN test that measures antibody titers interfering with Spike-receptor binding. Clearly higher surrogate VN titers were induced by the Spike^Opt^ antigen compared to the Spike^WT^ antigen (Figure 2B). These titers were boosted after the second vaccination with high titers until the end of the experiment. Consistently, the titers induced by Spike^Opt^ antigen were higher in comparison to the RP vaccine producing the Spike^WT^ antigen (Figures 2C-D).

The VEEV RP vector platform is known for its efficient induction of both humoral as well as cellular responses [24]. To assess the level of cellular responses induced by the RP vaccine candidates, a third immunization was performed and seven days later lymphocytes were isolated for a lymphocyte stimulation test (LST). All isolated lymphocytes stimulated with ConA resulted in >80% proliferation titers. In contrast to the differences in humoral responses between the Spike^WT^ and Spike^Opt^ antigens, no differences were observed in levels of SARS-CoV-2 S1 specific T-cell differentiation (Figure 2E). To determine whether the humoral responses also resulted in mucosal immunity, tracheal swabs were taken at the end of the experiments. Interestingly, also surrogate VN titers were detected in the trachea swabs, and the levels correlated with the systemic antibody levels with superior titers for the Spike^Opt^ antigen compared to the Spike^WT^ antigen (Figure 2F). These antibody titers suggest that parental vaccination could induce protective mucosal immunity.

**Figure 2:**
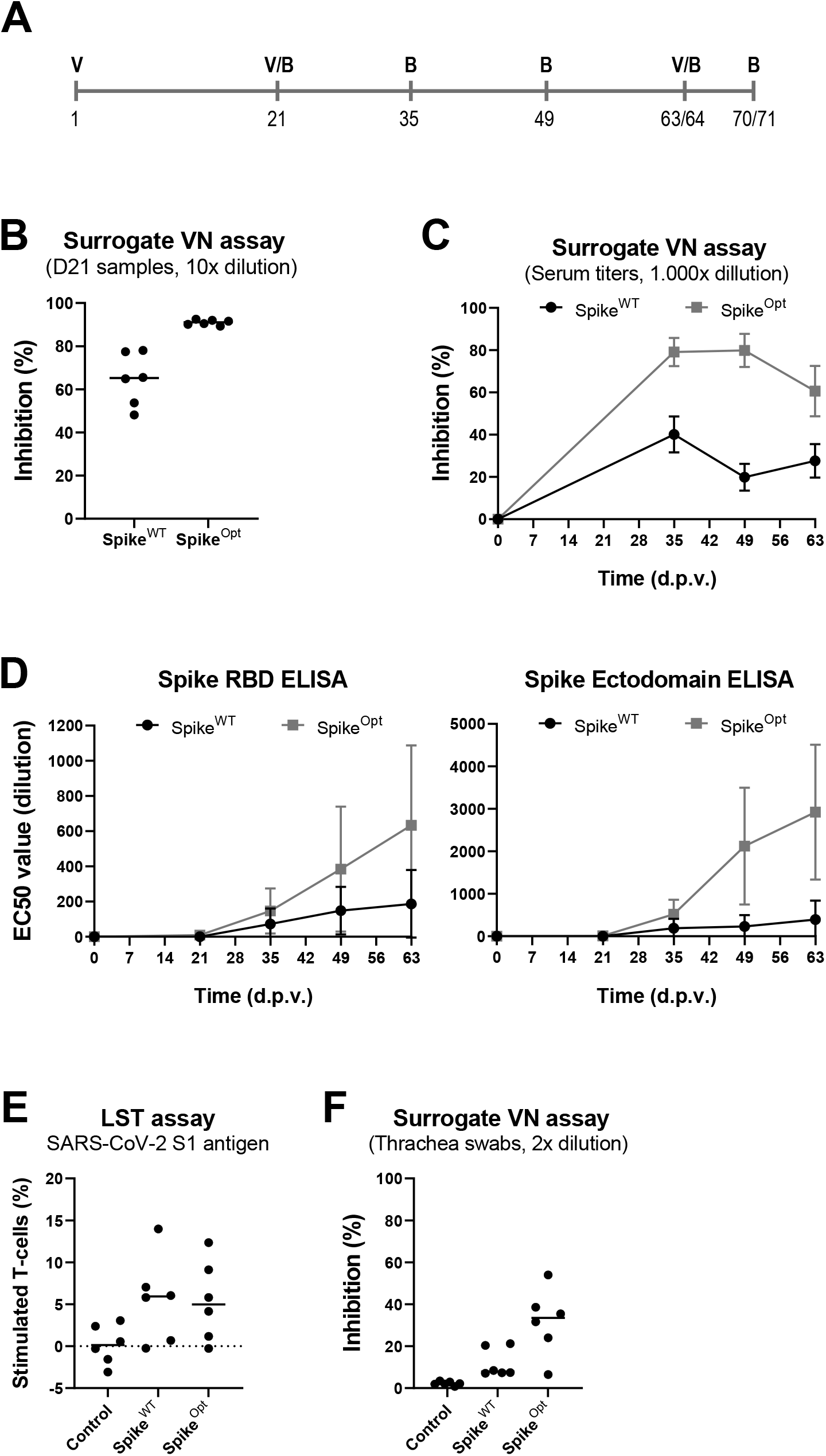
Immunogenicity study of vaccine candidates in a guinea pig model. (A) Overview of animal handlings. V = vaccination and B = blood sampling. (B) Surrogate SARS-CoV-2 virus neutralization (VN) test performed using 10-fold diluted serum samples from day 21 (D21). (C) Surrogate SARS-CoV-2 virus neutralization (VN) test performed using 1.000-fold diluted serum samples from day 35, 49 and 63/64 post prime vaccination (d.p.v.). Black line shows the antibody levels induced by the Spike^WT^ antigen and grey line shows the antibody levels induced by the Spike^Opt^ antigen. (D) Indirect ELISA results using the SARS-CoV-2 Spike RBD (left) or ectodomain (right) as antigen. Shown are EC50 values of sera (expressed as fold dilution) from cats exposed to the Spike^WT^ antigen (black line) or the Spike^Opt^ antigen (grey line). (E) Results of lymphocyte stimulation test (LST) from blood collected on day 70/71. Purified SARS-CoV-2 S1 antigen was used to stimulate isolated lymphocytes and proliferation was measured 96h after stimulation. (F) Surrogate VN test performed using 2-fold diluted swab samples taken at day 70/71.

### Cat vaccination-challenge study

To determine vaccine efficacy, cats were either vaccinated with a RP vaccine producing EGFP (Control), the optimized SARS-CoV-2 Spike antigen (Spike^Opt^) or remained non-vaccinated (sentinels). Three weeks post booster vaccination, cats were exposed to a mucosal SARS-CoV-2 challenge and samples were taken as outlined in Figure 3A.

**Figure 3:**
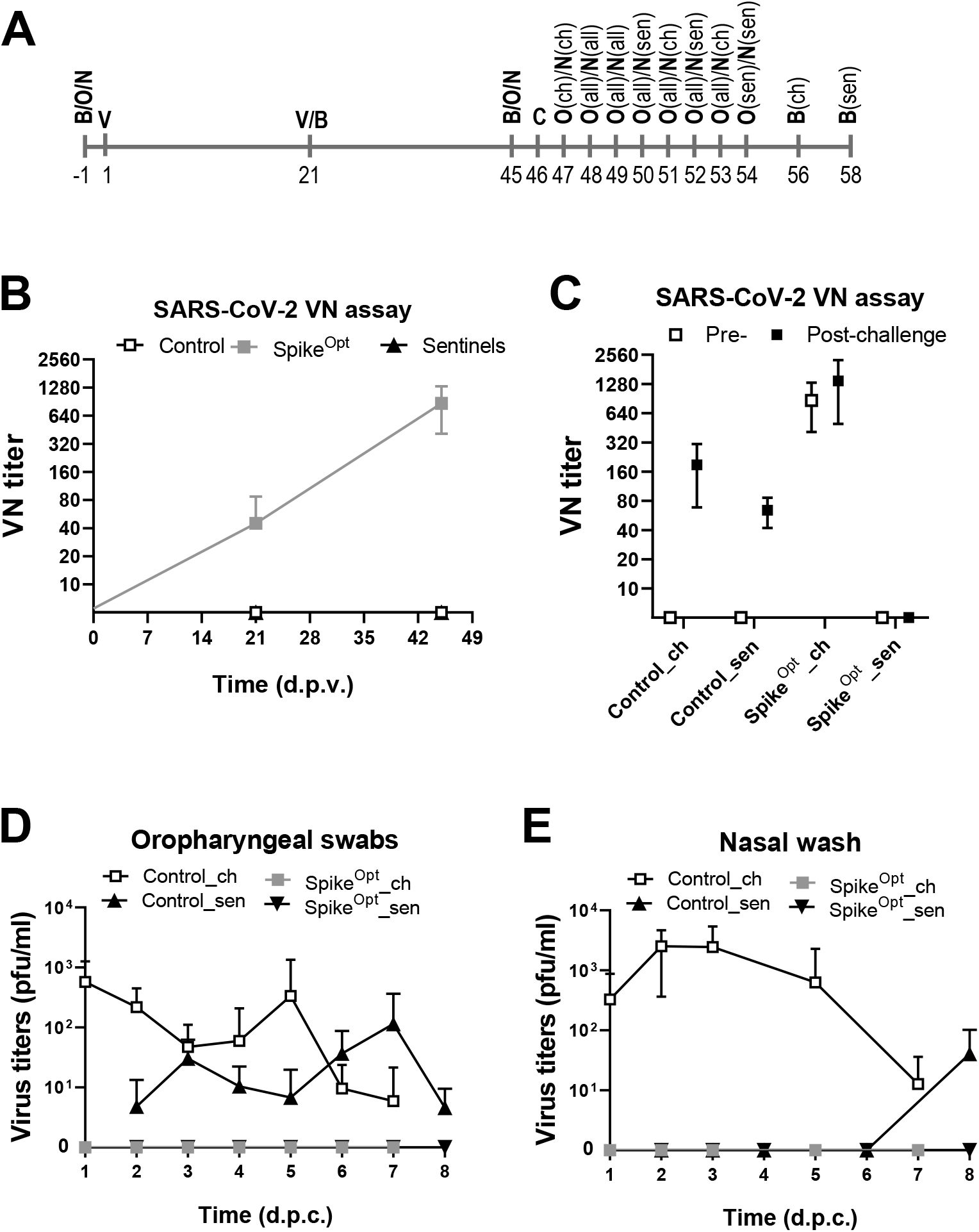
Vaccination-challenge experiment in cat (A) Overview of animal handlings. V = vaccination, B = blood sampling, O = oropharyngeal swabs, N = nasal wash, (all) = all animals, (ch) = only challenged animals, (sen) = only sentinel animals. (B) Serum neutralizing antibody titers determined using a SARS-CoV-2 virus neutralization (VN) test 21- and 45-days post vaccination (d.p.v.). Black line with open squares shows the antibody levels in the control vaccinated animals, black line with black triangles shows the antibody levels in non-vaccinated sentinel animal, and grey line with grey squares show antibody titers induced by the Spike^Opt^ antigen. (C) Serum neutralizing antibody titers determined using a SARS-CoV-2 virus neutralization (VN) test at day of challenge, 45-days post vaccination (open squares) and 12 (challenged) or 14 (sentinel) days post challenge (black squares). (D) SARS-CoV-2 virus titers in pfu/ml in oropharyngeal swabs 1 till 8 days post challenge (d.p.c.). Black line with open squares shows viral titers in challenged control animals, black line with black triangles shows viral titers in non-vaccinated sentinel animal co-housed with control animals, grey line with grey squared show viral titers in Spike^Opt^ antigen vaccinated animals, and black line with black inverted triangles show viral titers in non-vaccinated sentinel animals co-housed with Spike^Opt^ antigen vaccinated animals. (E) SARS-CoV-2 virus titers in pfu/ml in nasal wash after challenge. Lines and symbols as in (D).

Following vaccination, no adverse reactions were detected in any of the cats at any timepoint. The RP vaccine producing the Spike^Opt^ antigen was able to induce a virus neutralising antibody titer in all cats after a single vaccination, which was boosted after the second vaccination and maintained levels until the challenge 3.5 weeks later (Figure 3B). Control and non-vaccinated sentinel animals remained negative at all times up until challenge. Both the challenged and sentinel cats did not demonstrate any clinical signs post challenge. However, nine out of ten control challenged cats shed virus orally (Figure 3D) and nasally (Figure 3E) one day after challenge and for at least 3 days during the observation period. These data show that the mucosal SARS-CoV-2 challenge results in efficient virus replication in the respiratory tract. Higher and more consistent virus shed was detected from the nasal washes whereas the oropharyngeal swabs demonstrated a less consistent pattern, the reason for this is unknown. Interestingly, virus shed was also detected from the nasal washes in two of the non-vaccinated sentinels placed with the control animals one day after challenge. Moreover, all five sentinel animals shed virus via the oral route for at least two days demonstrating efficient spread of the virus from control challenged to sentinel animals (Figure 3D).

None of the vaccinated cats shed any detectable virus orally (Figure 3D) or nasally (Figure 3E) at any timepoint after the challenge. The results suggest that the vaccine may have prevented infection. Also, no virus was detected in the non-vaccinated sentinels housed with the vaccinated cats as would be expected considering the lack of challenge virus replication in the vaccinated cats. Analysis of virus neutralising antibody titers post challenge confirmed the findings that both control challenged and sentinel animals were efficiently infected (Figure 3C). In contrast, no seroconversion was observed in the sentinel animals housed with the vaccinated cats. Thus, the VEEV RP vaccine producing the Spike^Opt^ antigen appears to induce sterile immunity and prevent transmission from infected to naïve cats.

## Discussion

With the ongoing pandemic and reports of naturally occurring SARS-CoV-2 infections of a variety of animal species, it is important to understand the epidemiology of this virus in these animal populations especially with regards to the establishment of potential reservoirs, mutations and transmission within and to other species. SARS-CoV-2 infections in humans can be transmitted to cats and it has been hypothesised that cat to cat transmission of virus can take place in a natural setting [1]. It was previously demonstrated that SARS-CoV could infect and spread between cats [22] but the complete epidemiological picture of feline infection was not fully understood with the rapid eradication of SARS-CoV from humans before it reached a pandemic situation. The situation with SARS-CoV-2 is different as it has become a global issue with the likelihood of becoming endemic in the human population. Whilst an infected cat is considered to be low risk for SARS-CoV-2 transmission to humans, to other cats and other species, the fact that infected cats shed virus for prolonged periods which can potentially be aerosolised gives credence to the possibility that cats may play some role in the viral epidemiology either by transmitting the virus onwards, enabling further mutation, or acting as a virus reservoir. Although routine vaccination of cats is not proposed, should the epidemiological situation change, the availability of a vaccine which can be rapidly produced, updated, and which reduces or prevents viral replication and transmission between cats and other animals will be useful. Furthermore, a vaccine that could be used in a range of susceptible animal species would be preferable.

We have demonstrated that optimal expression of coronavirus Spike antigens is critical to the induction of a sustained neutralising antibody response in both guinea pigs and cats. In the guinea pig experiments it was interesting to note that intramuscular vaccination induced some level of mucosal antibody titers, which was somewhat surprising and is likely to be a wash over from serological induction. It remains to be established whether these antibodies might contribute to the sterile immunity that has been observed for this vaccine candidate in cats.

The optimised Spike RP vaccine successfully induced a virus neutralising antibody response in all vaccinated cats after a single vaccination which was boosted upon second vaccination. Furthermore, the vaccine was able to prevent infection in all vaccinated cats as demonstrated by the lack of virus re-isolation post challenge. Although there was a strong induction of a serological response in the cats it was not investigated, as was demonstrated in the guinea pig experiments, whether neutralising antibody was present in the respiratory tract. Furthermore, we did not examine the role of cell mediated immunity in the prevention of infection of cats nor were we able to extend the experiments to investigate the duration of the immune response induced. The ability to induce local protection from parenterally administered coronavirus vaccines is not well established and in certain veterinary respiratory coronavirus infections mucosally-applied live attenuated vaccines are used to reduce the viral replication at the site of initial infection. These live vaccines induce a relatively brief period of protection, so they are boosted by inactivated adjuvanted whole virus vaccines to establish longer immunity. Use of inactivated vaccines alone in these veterinary settings is not as effective at protecting the local respiratory tract as the live priming inactivated boost approach [23]. For this reason, it is reassuring that a parenterally administered vaccine did appear to provide respiratory protection in the feline model. Future work would be needed to establish whether a single vaccine dose would also be sufficient to protect the cats from infection. Furthermore, the optimal inoculation schedule has not yet been established for this vaccine nor importantly has the duration of immunity that can be induced and whether the ability to prevent infection and spread persists over this time.

This work demonstrates the utility of the VEEV strain TC-83-based replicon RNA particle vaccine platform (Sequivity^®^). RP vaccines based on VEEV have previously been shown to protect cats against viral diseases including some respiratory protection against clinical signs and virus shed in a feline calicivirus infection model, and in addition have been shown to be effective in multiple species including dogs, horses, pigs, cattle, chickens and ducks [24], authors unpublished observations]. Furthermore, VEEV based RP vaccines expressing the Spike proteins from other coronaviruses have been shown to induce virus neutralising antibodies [34]. The advantages of RP based technology is that vaccines can be rapidly prepared if the gene of interest is known to encode a protective antigen. This vaccine platform is safe-by-design as the RP vaccines undergo a non-productive cycle of replication in which replicon RNA but no virus is replicated and no adjuvants are required [27]. Thus far Rhesus macaques, hamsters and ferrets have been utilised as natural animal models for SARS-CoV-2. In these animals, infection is usually asymptomatic or induces mild clinical disease. As SARS-CoV-2 also induces asymptomatic infections in cats this species may also provide a possibility to study disease transmission, especially via aerosols and vaccine design aimed at preventing initial infection in the respiratory tract. In some respects, infection of cats may mimic the majority of human infections which are asymptomatic.

This work demonstrates that a replicon-based vaccine expressing the stabilised SARS-CoV-2 Spike protein was able to induce high levels of virus neutralising antibodies in serum of vaccinated cats and that the induced response was able to prevent infection of the upper respiratory tract thereby preventing onward transmission to other cats.

## Acknowledgements

We thank Michelle Allen for her assistance in conducting the animal experiments in guinea pigs. We thank Amber King, Danielle Egan, Rebecca Gillaspie, Kari Carritt, Joel Schrader, Eva Restis, Josette van den Berg, Maikel Schaap, and Pasqualina Fleuren for animal husbandry.

## References

[1] Shi J, Wen Z, Zhong G, Yang H, Wang C, Huang B, et al. Susceptibility of ferrets, cats, dogs, and other domesticated animals to SARS-coronavirus 2. Science 2020;1020:1016–20. https://doi.org/10.1126/science.abb7015.

[2] OIE Report n.d. https://www.oie.int/scientific-expertise/specific-information-and-recommendations/questions-and-answers-on-2019novel-coronavirus/events-in-animals/.

[3] Munnink BBO, Sikkema RS, Nieuwenhuijse DF, Molenaar RJ, Munger E, Molenkamp R, et al. Transmission of SARS-CoV-2 on mink farms between humans and mink and back to humans. Science (80-) 2021;371:172–7. https://doi.org/10.1126/science.abe5901.

[4] Bosco-Lauth AM, Hartwig AE, Porter SM, Gordy PW, Nehring M, Byas AD, et al. Experimental infection of domestic dogs and cats with SARS-CoV-2: Pathogenesis, transmission, and response to reexposure in cats. Proc Natl Acad Sci U S A 2020;117:26382–8. https://doi.org/10.1073/pnas.2013102117.

[5] Halfmann PJ, Hatta M, Chiba S, Maemura T, Fan S, Takeda M, et al. Transmission of SARS-CoV-2 in Domestic Cats. N Engl J Med 2020:8–9. https://doi.org/10.1056/nejmc2013400.

[6] Hosie MJ, Hofmann-Lehmann R, Hartmann K, Egberink H, Truyen U, Addie DD, et al. Anthropogenic Infection of Cats during the 2020 COVID-19 Pandemic. Viruses 2021;13. https://doi.org/10.3390/v13020185.

[7] Zhang Q, Zhang H, Gao J, Huang K, Yang Y, Hui X, et al. A serological survey of SARS-CoV-2 in cat in Wuhan. Emerg Microbes Infect 2020;9:2013–9. https://doi.org/10.1080/22221751.2020.1817796.

[8] Hamer SA, Pauvolid-Corrêa A, Zecca IB, Davila E, Auckland LD, Roundy CM, et al. Natural SARS-CoV-2 infections, including virus isolation, among serially tested cats and dogs in households with confirmed human COVID-19 cases in Texas, USA. BioRxiv Prepr Serv Biol 2020. https://doi.org/10.1101/2020.12.08.416339.

[9] Fritz M, Rosolen B, Krafft E, Becquart P, Elguero E, Vratskikh O, et al. High prevalence of SARS-CoV-2 antibodies in pets from COVID-19+ households. One Heal 2021;11:0–4. https://doi.org/10.1016/j.onehlt.2020.100192.

[10] Patterson EI, Elia G, Grassi A, Giordano A, Desario C, Medardo M, et al. Evidence of exposure to SARS-CoV-2 in cats and dogs from households in Italy. Nat Commun 2020;11. https://doi.org/10.1038/s41467-020-20097-0.

[11] Huang Q, Zhan X, Zeng X-T. COVID-19 pandemic: stop panic abandonment of household pets. J Travel Med 2020;27. https://doi.org/10.1093/jtm/taaa046.

[12] OIE Report n.d. https://www.oie.int/fileadmin/Home/MM/A_Reporting_SARS-CoV-2_to_the_OIE.pdf.

[13] Montagutelli X, Prot M, Levillayer L, Baquero Salazar E, Jouvion G, Conquet L, et al. The B1.351 and P.1 variants extend SARS-CoV-2 host range to mice. BioRxiv 2021:2021.03.18.436013. https://doi.org/10.1101/2021.03.18.436013.

[14] Ferasin L, Fritz M, Ferasin H, Becquart P, Legros V, Leroy EM. Myocarditis in naturally infected pets with the British variant of COVID-19. BioRxiv 2021:2021.03.18.435945. https://doi.org/10.1101/2021.03.18.435945.

[15] Krammer F. SARS-CoV-2 vaccines in development. Nature 2020;586:516–27. https://doi.org/10.1038/s41586-020-2798-3.

[16] Vennema H, de Groot RJ, Harbour DA, Dalderup M, Gruffydd-Jones T, Horzinek MC, et al. Early death after feline infectious peritonitis virus challenge due to recombinant vaccinia virus immunization. J Virol 1990;64:1407–9. https://doi.org/10.1128/JVI.64.3.1407-1409.1990.

[17] Bande F, Arshad SS, Omar AR, Hair-Bejo M, Mahmuda A, Nair V. Global distributions and strain diversity of avian infectious bronchitis virus: A review. Anim Heal Res Rev 2017;18:70–83. https://doi.org/10.1017/S1466252317000044.

[18] Davelaar FG, Noordzij A, Van Der Donk JA. A study on the synthesis and secretion of immunoglobulins by the harderian gland of the fowl after eyedrop vaccination against infectious bronchitis at 1-day-old. Avian Pathol 1982;11:63–79. https://doi.org/10.1080/03079458208436082.

[19] Toro H, Fernandez I. Avian infectious bronchitis: specific lachrymal IgA level and resistance against challenge. Zentralbl Veterinarmed B 1994;41:467–72. https://doi.org/10.1111/j.1439-0450.1994.tb00252.x.

[20] Orr-Burks N, Gulley SL, Toro H, van Ginkel FW. Immunoglobulin A as an early humoral responder after mucosal avian coronavirus vaccination. Avian Dis 2014;58:279–86. https://doi.org/10.1637/10740-120313-Reg.1.

[21] Cavanagh D, Darbyshire JH, Davis P, Peters RW. Induction of humoral neutralising and haemagglutination-inhibiting antibody by the spike protein of avian infectious bronchitis virus. Avian Pathol 1984;13:573–83. https://doi.org/10.1080/03079458408418556.

[22] Song CS, Lee YJ, Lee CW, Sung HW, Kim JH, Mo IP, et al. Induction of protective immunity in chickens vaccinated with infectious bronchitis virus S1 glycoprotein expressed by a recombinant baculovirus. J Gen Virol 1998;79 (Pt 4):719–23. https://doi.org/10.1099/0022-1317-79-4-719.

[23] Ignjatovic J, Galli L. The S1 glycoprotein but not the N or M proteins of avian infectious bronchitis virus induces protection in vaccinated chickens. Arch Virol 1994;138:117–34. https://doi.org/10.1007/BF01310043.

[24] Mogler MA, Kamrud KI. RNA-based viral vectors. Expert Rev Vaccines 2015;14:283–312. https://doi.org/10.1586/14760584.2015.979798.

[25] Vander Veen RL, Loynachan AT, Mogler MA, Russell BJ, Harris DLH, Kamrud KI. Safety, immunogenicity, and efficacy of an alphavirus replicon-based swine influenza virus hemagglutinin vaccine. Vaccine 2012;30:1944–50. https://doi.org/10.1016/j.vaccine.2012.01.030.

[26] Pushko P, Parker M, Ludwig G V, Davis NL, Johnston RE, Smith JF. Replicon-helper systems from attenuated Venezuelan equine encephalitis virus: expression of heterologous genes in vitro and immunization against heterologous pathogens in vivo. Virology 1997;239:389–401. https://doi.org/10.1006/viro.1997.8878.

[27] Kamrud KI, Alterson K, Custer M, Dudek J, Goodman C, Owens G, et al. Development and characterization of promoterless helper RNAs for the production of alphavirus replicon particle. J Gen Virol 2010;91:1723–7. https://doi.org/10.1099/vir.0.020081-0.

[28] Amanat F, Strohmeier S, Rathnasinghe R, Schotsaert M, Coughlan L, García-Sastre A, et al. Introduction of two prolines and removal of the polybasic cleavage site leads to optimal efficacy of a recombinant spike based SARS-CoV-2 vaccine in the mouse model. BioRxiv 2020;2:2020.09.16.300970. https://doi.org/10.1101/2020.09.16.300970.

[29] Mercado NB, Zahn R, Wegmann F, Loos C, Chandrashekar A, Yu J, et al. Single-shot Ad26 vaccine protects against SARS-CoV-2 in rhesus macaques. Nature 2020. https://doi.org/10.1038/s41586-020-2607-z.

[30] Hsieh C-L, Goldsmith JA, Schaub JM, DiVenere AM, Kuo H-C, Javanmardi K, et al. Structure-based design of prefusion-stabilized SARS-CoV-2 spikes. Science 2020;369:1501–5. https://doi.org/10.1126/science.abd0826.

[31] Hooper JW, Ferro AM, Golden JW, Silvera P, Dudek J, Alterson K, et al. Molecular smallpox vaccine delivered by alphavirus replicons elicits protective immunity in mice and non-human primates. Vaccine 2009;28:494–511. https://doi.org/10.1016/j.vaccine.2009.09.133.

[32] Kamrud KI, Custer M, Dudek JM, Owens G, Alterson KD, Lee JS, et al. Alphavirus replicon approach to promoterless analysis of IRES elements. Virology 2007;360:376–87. https://doi.org/10.1016/j.virol.2006.10.049.

[33] Sanders RW, Moore JP. Virus vaccines: proteins prefer prolines. Cell Host Microbe 2021;29:327–33. https://doi.org/10.1016/j.chom.2021.02.002.

[34] Sawattrakool K, Stott CJ, Bandalaria-Marca RD, Srijangwad A, Palabrica DJ, Nilubol D. Field trials evaluating the efficacy of porcine epidemic diarrhea vaccine, RNA (Harrisvaccine) in the Philippines. Trop Anim Health Prod 2020;52:2743–7. https://doi.org/10.1007/s11250-020-02270-1.

